# Demyelination induces selective vulnerability of inhibitory networks in multiple sclerosis

**DOI:** 10.1101/2020.09.17.302042

**Authors:** Lida Zoupi, Sam A. Booker, Dimitri Eigel, Carsten Werner, Peter C. Kind, Tara L. Spires-Jones, Ben Newland, Anna C. Williams

## Abstract

In multiple sclerosis (MS), a chronic demyelinating disease of the central nervous system, neurodegeneration is detected early in the disease course and is associated with the long-term disability of patients. Neurodegeneration is linked to both inflammation and demyelination, but its exact cause remains unknown. This gap in knowledge contributes to the current lack of treatments for the neurodegenerative phase of MS. Here we ask if neurodegeneration in MS affects specific neuronal components and if it is the result of demyelination. Neuropathological examination of secondary progressive MS motor cortices revealed a selective vulnerability of inhibitory interneurons in MS. The generation of a rodent model of focal subpial cortical demyelination proved that this selective neurodegeneration is secondary to demyelination providing the first temporal evidence of demyelination-induced neurodegeneration and a new preclinical model for the study of neuroprotective treatments.

## Introduction

Multiple sclerosis (MS) is a chronic neuroinflammatory disease of the human central nervous system (CNS) characterised by inflammation, focal areas of demyelination and neurodegeneration. At the early stages of the disease, inflammatory cells target the myelin sheath that insulates and supports the axons. Current treatments that modulate this immune attack suppress the inflammatory demyelinating white matter lesions (as assessed by magnetic resonance imaging MRI) and accompanying relapses, but ultimately fail to prevent neurodegeneration (1).

Neurodegeneration in MS is reported to consist of neurite transection, diffuse synaptic dysfunction and overall neuronal loss (2–7). Long term disability better correlates with brain atrophy on MRI rather than with white matter demyelinating lesion load, but the mechanisms leading to these changes remain largely unknown (8–10). This limited understanding may contribute to the lack of successful therapies counteracting neurodegeneration. Thus, two fundamental questions need to be addressed: first, which neural components are primarily affected? Second, does neurodegeneration result directly from demyelination?

Given that cortical thinning/atrophy in MS is widespread, one may predict that all neuronal classes are potentially affected. However, in a recent transcranial magnetic stimulation study, motor dysfunction strongly correlates with increased excitatory and reduced inhibitory transmission in the motor cortex of MS patients, suggesting possible differential effects (11). Although demyelination in MS is classically identified in the white matter, due to the relative ease of identification on MRI scans, it is well-recognised that demyelination is also common in the grey matter (12–15). Focal demyelinated lesions occur in the grey matter, including in the cortex, disrupting neuronal signaling directly or via an inflammatory toxic environment (16, 17). Differential effects of these lesions on various cortical neurons is possible due to spatial patterning, with lesions identified in both superficial (subpial lesions) and deeper cortical layers (intracortical, leukocortical lesions) (12, 13, 15). Furthermore, the pattern of myelination of neuronal subtypes is variable, potentially increasing their susceptibility to neurodegeneration via demyelination: only some subtypes of cortical inhibitory interneurons are myelinated and while some pyramidal excitatory neuronal axons are fully myelinated along their length, others have discontinuous patterns of myelination, with large unmyelinated gaps between myelin sheaths (18–23).

Notably, it is still controversial as to whether or to what extent neurodegeneration is caused by demyelination directly. Proponents indict the evidence that demyelination deprives the axon of insulation and essential metabolic supplies from the oligodendrocyte which can lead to axonal degeneration in animal models, via mitochondrial dysfunction and persistent oxidative stress (24, 25). Supporting this, in chronic inactive demyelinated MS lesions, which by definition have little inflammation, there is slow axonal disintegration (2, 26). However, opponents of this theory point out that demyelination in MS is also accompanied by (or driven by) inflammation, including cytotoxic T cells and macrophages, which can trigger acute axonal damage by releasing neurotoxic factors such as cytokines and reactive oxygen species outside the areas of demyelination and perhaps unconnected to it (16, 17, 27–29). Supporting this, is the widespread neurodegeneration seen in MS, which does not correlate with the number of demyelinated MS lesions at least in the white matter (9). Furthermore, in the rodent model of experimental autoimmune encephalomyelitis (EAE), inflammation appears to mediate demyelination and neurodegeneration independently (5).

A better understanding of what neurodegeneration consists of in MS, and the mechanism of damage, may aid the development of therapeutic strategies to limit it. To address this, we first asked if there is specific vulnerability of different neurons to neurodegeneration in MS brains. By neuropathological examination of the motor cortex of secondary progressive MS patients, compared to non-neurological controls, we found that inhibitory neurons are more susceptible to neurodegeneration. Next, to examine if grey matter demyelination directly causes neurodegeneration, we generated a rodent model of focal cortical subpial demyelination that also shows selective inhibitory neuron and synapse loss and provided temporal evidence of demyelination-induced neurodegeneration. Our data support the idea that neurodegeneration can be secondary to demyelination irrespective of the presence of diffuse inflammation in the cortex, whilst developing a preclinical model for screening drugs to improve neuroprotection in MS patients.

## Results

### Selective loss of myelinated interneurons and inhibitory synapses in the MS motor cortex

To assess the effects of MS on cortical structure, we first examined gross neurodegeneration in the motor cortex of post-mortem secondary progressive MS tissue and non-neurological controls. Grey matter (GM) thickness, measured as the distance between layers 1 to 6, was similar between control and MS samples **(Supplementary Fig. 1A)**. However, there was significant axonal loss in the MS motor cortex, as quantified using the pan-axonal marker SMI312 **(Supplementary Fig. 1B)**, in accordance with studies reporting broad axonal loss in MS GM.

By their location in layers 2/3, these lost axons are either cortico-cortical projection axons, excitatory axon collaterals or inhibitory interneuron axons. Therefore, we next checked for changes in density of excitatory and inhibitory synapses between control and MS motor cortices, assuming overlap of pre- and post-synaptic puncta as evidence of a functional synapse. Using confocal microscopy and array tomography, we stained sections for the pre-synaptic marker synapsin1 (SYN1) and the excitatory post-synaptic marker post-synaptic density protein 95 (PSD95) **(Fig. 1A,B, Supplementary Fig. 2A-C)** or the inhibitory pre-synaptic marker vesicular GABA transporter (VGAT) and post-synaptic marker GEPHYRIN **(Fig. 1D,E)**. Synapses were quantified in layers 2/3 of the cortex where the majority of cortico-cortical connections are formed, using automated methods previously described (30). Both methods showed that there is no difference in the density of excitatory synapses in the MS motor cortex when compared to controls **(Fig. 1C & Supplementary Fig. 2D)**. However, we observed a significant 25% reduction in inhibitory synapses in MS motor cortices compared to controls **(Fig. 1F)**.

**Fig. 1.**
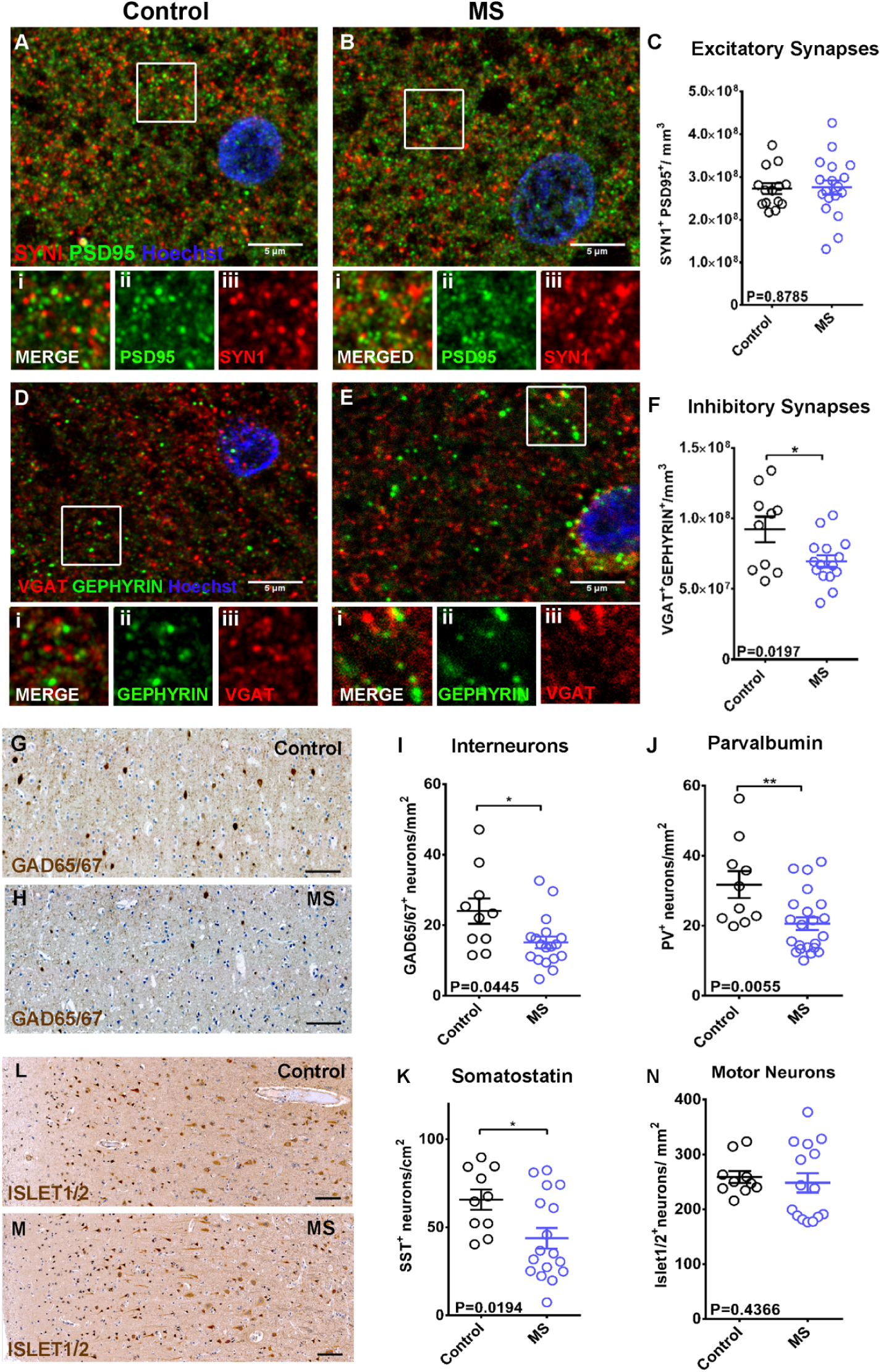
**A-B:** Immunohistochemistry of control (A) and MS (B) motor cortex (layers 2/3) for SYN1 (red), PSD95 (green), with Hoechst (blue). Scale bar: 5µm. Inset images of boxed areas in A or B showing merged (i) and single channels of the synaptic proteins PSD95 (ii) and SYN1 (iii). **C:** Quantification of SYN1+/PSD95+ excitatory synapses in layers 2/3 neuropil of control and MS motor cortex (Control: mean 2.723e+008 ± 1.244e+007 SEM synapses/mm^3^, MS: mean 2.757e+008 ± 1.665e+007SEM synapses/mm^3^; each point is a patient, unpaired t test).**D-E:** Immunohistochemistry of control (D) and MS (E) motor cortex (layers 2/3) for VGAT (red) GEPHYRIN (green) with Hoechst (blue). Scale bar: 5µm. Inset images of boxed areas in D or E showing merged (i) and single channels of the synaptic proteins GEPHYRIN (ii) and VGAT (iii). **F:** Quantification of VGAT+/GEPHYRIN+ inhibitory synapses in layers 2/3 neuropil of control and MS motor cortex (Control: mean 9.217e+007 ± 8.997e+006 SEM synapses/mm^3^, MS: mean 6.966e+007 ± 4.272e+006 SEM synapses/mm^3^; each point is a patient, unpaired t test). **G-H:** Immunohistochemistry of human control (G) and MS (H) motor cortex for the pan-interneuron marker GAD65/67 (brown) with hematoxylin counterstain (blue). Scale bar: 100µm. **I:** Quantification of GAD65/67+ interneuron density in all six cortical layers in control and MS motor cortex (Control: mean 23.99 ± 3.615 SEM neurons/mm^2^, MS: mean 15.13 ± 1.675 SEM neurons/mm^2^; each point is a patient, unpaired t test). **J:** Quantification of PV+ interneuron density in all six cortical layers in control and MS motor cortex (Control: mean 31.74 ± 3.834 SEM neurons/mm^2^, MS: mean 20.60 ± 1.829 SEM neurons/mm^2^; each point is a patient, unpaired t test). **K:** Quantification of SST+ interneuron density in all six cortical layers in control and MS motor cortex (Control: mean 65.68 ± 5.713 SEM neurons/cm^2^, MS: mean 43.75 ± 5.809 SEM neurons/cm^2^; each point is a patient, unpaired t test). **L-M:** Immunohistochemistry of human control (L) and MS (M) motor cortex for the motor neuron marker Islet1/2 (brown) with hematoxylin counterstain (blue). Scale bar: 100µm. **N:** Quantification of Islet1/2+ motor neuron density in all six cortical layers in control and MS motor cortex (Control: 258.6±10.95SEM neurons/mm^2^, MS: mean 248.1±17.75SEM neurons/mm^2^; each point is a patient, Mann Whitney test).

We then asked if these changes in inhibitory synapse density can be attributed to a selective loss of inhibitory interneurons, using an antibody, raised against glutamate decarboxylases 65 and 67 (GAD65/67) that identifies all interneurons. We observed a significant reduction in the density of GAD65/67 positive neurons in MS motor cortex when compared to controls **(Fig. 1G,H,I)**. As not all interneurons are myelinated, we investigated whether there was a differential effect on interneuron subtypes. To do this, we immunolabelled cortical slices using antibodies against the calcium binding proteins parvalbumin (PV), calretinin (CR), calbindin (CB), or the neuropeptides somatostatin (SST) and cholecystokinin (CCK). We found a significant reduction in the density of PV expressing (PV+) and SST-expressing (SST+) interneurons (which usually have myelinated axons) **(Fig. 1J.K)**, but no change in the density of CRL, CCK, and CB expressing subtypes, which typically have unmyelinated axons **(Supplementary Fig. 1C-E)**. In accordance with the lack of change of excitatory synaptic densities, there was no difference in the densities of excitatory motor neurons (using an antibody against the LIM-homeobox domain markers Islet1/2) **(Fig. 1L, M,N)** between control and MS cortex indicating that they survive even in late progressive MS.

This selective reduction of two interneuron subtypes, PV+ and SST+ ones, which are normally myelinated, suggested an increased susceptibility of these to neurodegeneration in MS. These changes were present in areas of motor cortex that appeared to possess typical levels of myelination (normal appearing grey matter-NAGM) and those that were clearly demyelinated **(Supplementary Fig. 1F-H)**, with the caveat that NAGM areas show similar axonal loss to the demyelinated areas **(Supplementary Fig. 1I)** and so cannot be considered as normal healthy tissue. These results clearly show that specific interneuron subtypes are susceptible to neurodegeneration in the cortex of MS patients. Hence, we next tested whether cortical demyelination drives this pattern of neurodegeneration by developing an animal model.

### Loss of inhibitory neurons and synapses in mouse cortical demyelinated lesions

To investigate whether this selective neurodegeneration is driven by demyelination, we developed a novel, toxin-induced mouse model of focal subpial cortical demyelination. Using a macroporous poly(ethylene glycol) (PEG) based cylindrical scaffold (termed cryogel), we delivered the demyelinating toxin lysophosphatidylcholine (LPC) directly onto the surface of the motor cortex **(Fig. 2A)**. We have previously shown that this method generates focal cortical lesions ex vivo, in organotypic slice cultures (31). In vivo, a single subpial focal demyelinating lesion was formed below the LPC-loaded cryogel at two weeks post-surgery **(Fig. 2E,H)** as shown by the lack of myelin basic protein (MBP) signal at the lesion site and reduced signal in the perilesion area in the upper cortical layers (defined as the area extending 150 µm from the lesion border in cortical layers 2/3 **Supplementary Fig. 3A**). In both lesion and perilesion sites, there were increased numbers of ionized calcium binding adaptor molecule 1 (IBA1)-positive microglia/macrophages **(Supplementary Fig. 3C,G)** and glial fibrillary acidic protein (GFAP)-positive astrogliosis **(Supplementary Fig. 3H,I,L)**.

**Fig. 2.**
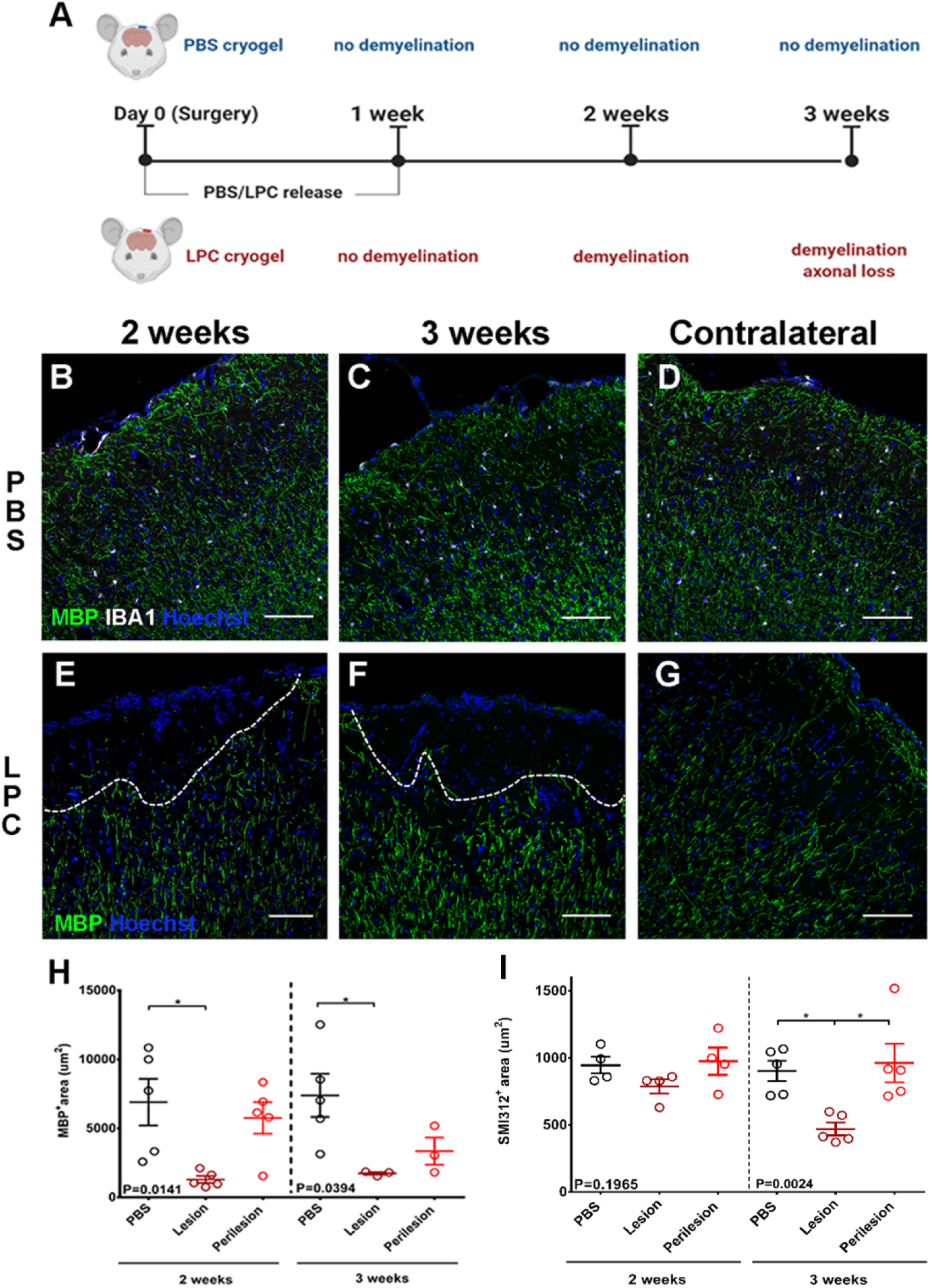
**A.** Outline of the experimental design. **B-D:** Immunohistochemistry of PBS cryogel treated animals for MBP (green), IBA1 (white) with Hoechst (blue), two (B) and three weeks (C) after cryogel placement. Contralateral hemisphere of the 2 week, PBS-loaded cryogel (D). Scale bar: 100µm. **E-G:** Immunohistochemistry of LPC cryogel treated animals for MBP (green) with Hoechst (blue) two (E) and three weeks (F) after cryogel placement (lesions outlined with dashed lines). Contralateral hemisphere of the 2 week, LPC-loaded cryogel (G). Scale bar: 100µm. **H:** Quantification of MBP+ signal at the upper cortical layers of PBS treated cortex (control) LPC-treated lesion and perilesion areas, two (left) and three (right) weeks post –surgery (2weeks PBS: mean 6900 ± 1692 SEM µm^2^, lesion: mean 1302 ± 250.2 SEM µm^2^, perilesion: mean 5756 ± 1139 SEM µm^2^; 3weeks PBS: mean 7388 ± 1561SEM µm^2^, lesion: mean 1733 ± 99.51 SEM µm^2^, perilesion: mean 3349 ± 984.8 SEM µm^2^ each point is an animal, One-way ANOVA Tukey’s multiple comparisons test). **I:** Quantification of SMI312+ signal at the upper cortical layers of PBS treated cortex (control) LPC-treated lesion and perilesion areas, two (left) and three (right) weeks post –surgery (2weeks PBS: mean 947.1± 63.17SEM µm^2^, lesion: mean 787.8± 52.70SEM µm^2^, perilesion: mean 976.3± 101.3SEM µm^2^; 3weeks PBS: mean 903.2 ± 75.53 SEM µm^2^, lesion: mean 471.1 ± 47.52 SEM µm^2^, perilesion: mean 962.9 ± 144.8 SEM µm^2^ each point is an animal, Kruskal-Wallis test with Dunn’s multiple comparisons test).

No effects on gliosis or myelin were observed in the contralateral hemisphere **(Fig. 2D,G,Supplementary Fig. 3B,D,F)** or when PBS-loaded cryogels were applied **(Fig. 2B,C & Supplementary Fig. 3H,J)**. At 3 weeks post-surgery, the focal demyelination persisted **(Fig. 2F,H)**, but with a significant reduction of microglia in the lesion compared to 2 weeks post-surgery **(Supplementary Fig. 3E,G)** and an attenuation of astrogliosis **(Supplementary Fig. 3J,K,L)**. Axonal loss was not marked until at 3 weeks post-surgery (approximately 50% reduction compared to controls) **(Fig. 2I)**. Very few CD3-expressing T-cells were seen in this model, restricted only to the area underneath the LPC-loaded cryogel at 2 weeks and subsiding by 3 weeks post-surgery **(Supplementary Fig. 3M-O)**.

Using our model, we asked if demyelination can directly lead to selective neurodegeneration. We found a reduced density of PV+ interneurons at 2 weeks post-surgery in the LPC treated motor cortex **(Fig. 3A-C)** and this persisted at 3 weeks post-surgery **(Fig. 3D)**. Additionally, we assessed damage to PV+ neurites (axons, dendrites) by quantifying the area covered by PV+ puncta, excluding PV+ cell bodies, **(Fig. 3E,F)** which revealed a marked loss in both lesion and perilesion areas by 3 weeks post-surgery **(Fig. 3G,H)**, accounting for the observed total axonal loss seen in **Fig. 2I**. In contrast, LPC-induced focal subpial demyelination did not alter the densities of vesicular glutamate transporter 2 (VGLUT2) expressing excitatory pyramidal neurons in cortical layers 2/3 when compared to PBS controls at 2 or 3 weeks post-surgery **(Fig. 3I-L)**; consistent with the human post-mortem data.

**Fig. 3.**
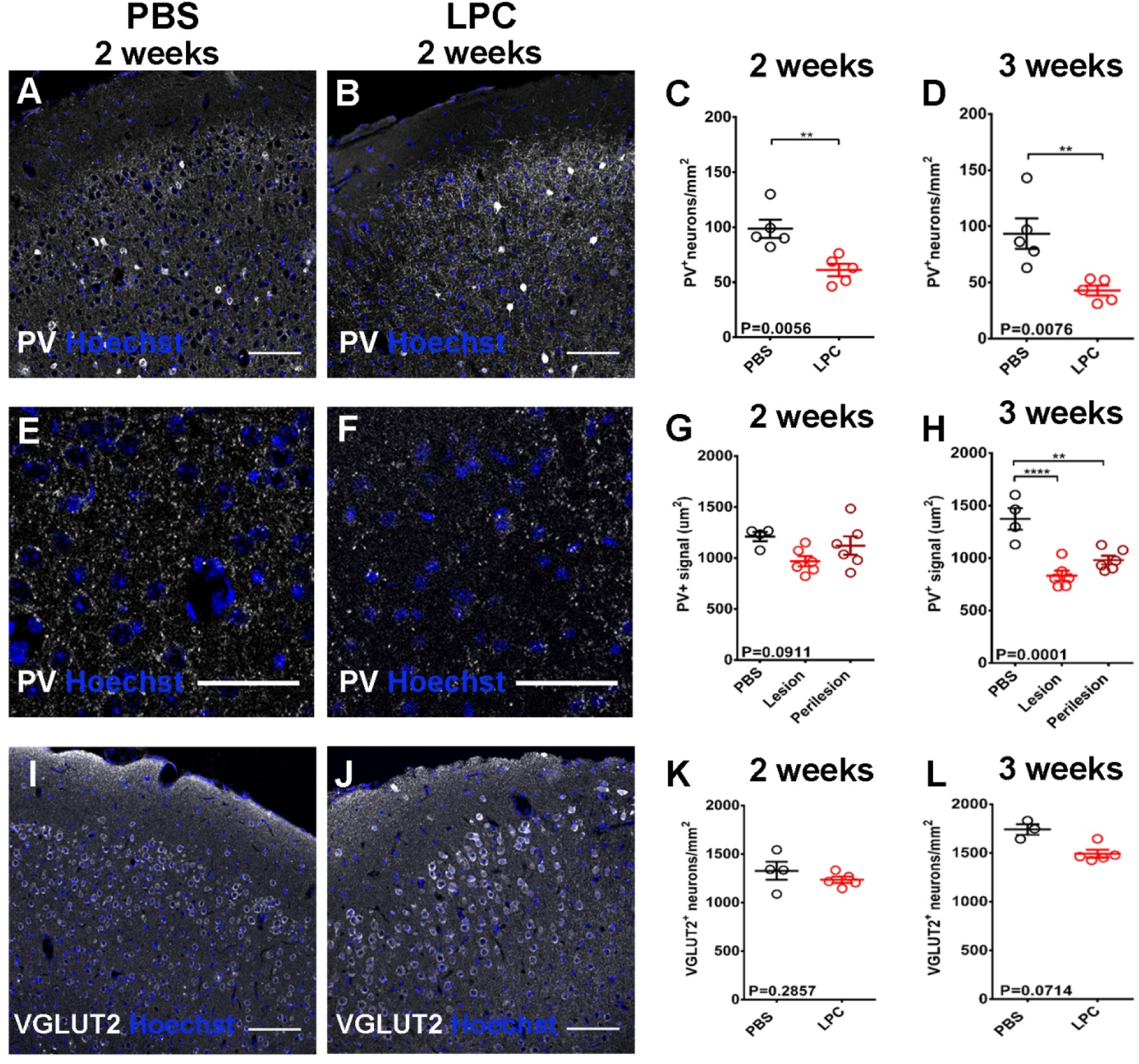
**A-B:** Immunohistochemistry of PBS (A) and LPC (B) treated animals for PV (white) with Hoechst (blue) two weeks post-surgery. Scale bar: 100µm. **C-D:** Quantification of PV+ neurons in layers 2/3 of PBS and LPC-treated animals two weeks (C, PBS: mean 98.63 ± 8.325 SEM neurons/mm^2^, LPC: mean 61.15 ± 5.520 SEM neurons/mm^2^; each point is an animal, unpaired t test) and three weeks post-surgery (D, PBS: mean 93.51 ± 13.58 SEM neurons/mm^2^, LPC: mean 42.89 ± 4.479 SEM neurons/mm^2^; each point is an animal, unpaired t test). **E-F:** Immunohistochemistry of PBS (E) and LPC (F) treated animals for PV (white) with Hoechst (blue) focusing on PV+ puncta at layers 2/3, two weeks post-surgery. Scale bar: 50µm. **G-H:** Quantification of PV+ area in layers 2/3 of PBS and LPC-treated animals two weeks (G, PBS: mean 1211 ± 45.12 SEM µm^2^, Lesion: mean 831.7 ± 47.26 SEM µm^2^, Perilesion: 981.0 ± 40.64 SEM µm^2^; each point is an animal, One-way ANOVA Tukey’s multiple comparisons test) and three weeks post-surgery (H, PBS: mean 1373 ± 102.4SEM µm^2^, Lesion: mean 968.1 ± 50.00SEM µm^2^, Perilesion: 1121 ± 90.13 SEM µm^2^; mean ± SEM, each point is an animal, One-way ANOVA Tukey’s multiple comparisons test). **I-J:** Immunohistochemistry of PBS (I) and LPC (J) treated animals for VGLUT2 (white) with Hoechst (blue) two weeks post-surgery. Scale bar: 100µm. **K-L:** Quantification of VGLUT2+ neurons in layers 2/3 of PBS and LPC-treated animals two weeks (K, PBS: mean 1328 ± 93.33 SEM neurons/mm^2^, LPC: mean 1234 ± 31.72 SEM neurons/mm^2^; each point is an animal, Mann Whitney test) and three weeks post-surgery (L, PBS: mean 1741 ± 54.65 SEM neurons/mm^2^, LPC: mean 1492 ± 39.51SEM neurons/mm^2^; each point is an animal, Mann Whitney test).

The impact of demyelination on the inhibitory network was further corroborated by a differential effect on the density of inhibitory **(Fig. 4A,B)** and excitatory synapses **(Fig. 4E,F)** between PBS control and LPC-lesioned animals at 2 and 3 weeks post-surgery. Fewer inhibitory synapses were detected in the demyelinated lesions by 3 weeks post-surgery compared to PBS controls as measured by automated quantification of the density of overlapping pre-synaptic VGAT and post-synaptic GEPHYRIN puncta **(Fig. 4D)**, with the reduction initiating at 2 weeks **(Fig. 4C)**. However, excitatory synapses (overlapping presynaptic vesicular glutamate transporter 1 (VGLUT1) and post-synaptic PSD95 from PSD95-GFP reporter transgenic mouse) did not differ between lesion and control cortex at either time point analysed **(Fig. 4 G,H)**.

**Fig. 4.**
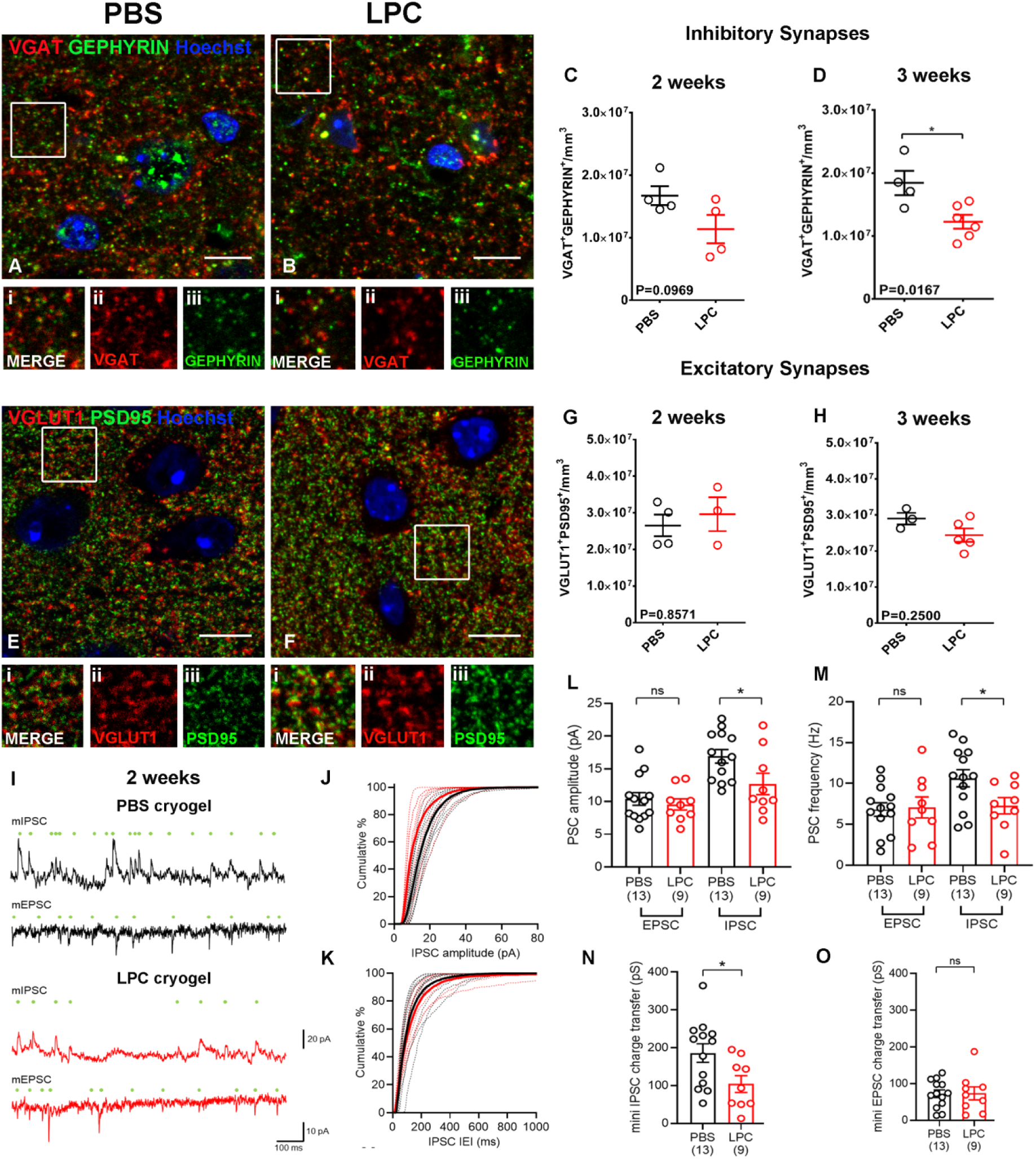
**A-B:** Immunohistochemistry of PBS (A) and LPC (B) treated animals for VGAT (red), GEPHYRIN (green) with Hoechst (blue) two weeks post-surgery. i-iii: Inset images of boxed areas in A or B showing merged (i) and single channels of the synaptic proteins VGAT (ii) and GEPHYRIN (iii). Scale bar: 10µm. **C-D:** Quantification of VGAT1+/GEPHYRIN+ inhibitory synapses in layers 2/3 of PBS and LPC-treated animals two (C, PBS: mean 1.673e+007 ± 1.520e+006SEM synapses/mm^3^, LPC: mean 1.138e+007 ± 2.255e+006 SEM synapses/mm^3^; each point is an animal, unpaired t test) and three weeks post-surgery (D, PBS: mean 1.844e+007 ± 1.936e+006 SEM synapses/mm^3^, LPC: mean 1.226e+007 ± 1.088e+006 SEM synapses/mm^3^; each point is an animal, unpaired t test). **E-F:** Immunohistochemistry of PBS (E) and LPC (F) treated animals for VGLUT1 (red), PSD95 (green) with Hoechst (blue) two weeks post-surgery. **i-iii:** Inset images of boxed areas in E or F showing merged (i) and single channels of the synaptic proteins VGLUT1 (ii) and PSD95 (iii). Scale bar: 10µm. **G-H:** Quantification of VGLUT1+/PSD95+ excitatory synapses in layers 2/3 of PBS and LPC-treated animals two weeks (G, PBS: mean 2.653e+007 ± 2.963e+006 SEM synapses/mm^3^, LPC: mean 2.961e+007 ± 4.605e+006 SEM synapses/mm^3^; each point is an animal, Mann Whitney test) and three weeks post-surgery (H, PBS: mean 2.899e+007 ± 1.597e+006 SEM synapses/mm^3^, LPC: mean 2.438e+007 ± 1.875e+006 SEM synapses/mm^3^; each point is an animal, Mann Whitney test). **I:** Representative miniature inhibitory postsynaptic currents (mIPSC) and miniature excitatory postsynaptic currents (mEPSC) recorded at 0 mV and −70 mV respectively in the presence of 500 nM TTX, from Layer 2 pyramidal cells from the motor cortex, two weeks post-surgery. Individual detected miniature events are indicated (green dots). Data are shown from representative cells from PBS (upper, black) and LPC (lower, red) treated mice. **J:** Cumulative distribution of mIPSC amplitudes from PBS (black) and LPC (red) treated mice, recorded at two weeks following surgery. Individual distributions from each neuron (thin dashed lines) are shown underlying the mean distribution (thick lines). **K:** Cumulative distributions of IPSC inter-event interval (IEI) recorded at two weeks post-surgery, plotted according to the same scheme as J. **L:** Amplitudes of mEPSC and mIPSCs recorded from Layer 2 pyramidal neurons in PBS (black) and LPC (red) treated mice at two weeks post-surgery (5 PBS mice and 3 LPC mice). Number of recorded neurons is indicated. **M:** Frequencies of mEPSC and mIPSCs recorded from layer 2 pyramidal neurons in PBS (black) and LPC (red) treated mice at two weeks post-surgery (5 PBS mice and 3 LPC mice). Number of recorded neurons is indicated. **N:** Total charge transferred by mIPSCs recorded at 0 mV for 5 minutes at two weeks post-surgery (5 PBS mice and 3 LPC mice). Number of recorded neurons is indicated. **O:** Total charge transferred by mEPSCs recorded at −70 mV over 5 minutes of recording from PBS and LPC treated mice at two weeks post-surgery (5 PBS mice and 3 LPC mice). Number of recorded neurons is indicated. Statistics shown: non-significance (ns) – *p* >.05, * *p* < .05, unpaired t-tests with Welch’s correction.

In summary, the generation of a single and focal subpial demyelinated cortical lesion in the mouse motor cortex caused a significant and early reduction in the numbers of PV+ interneuron somata and corresponding neurites and synapses, but without any change in excitatory neuron number and synaptic density. Our results successfully replicate the pattern of neurodegeneration seen in the human MS motor cortex and show that focal demyelination can directly lead to this neurodegeneration.

### Functional loss of inhibition is the earliest event after lesion formation

Histological evidence from our mouse model indicates a selective effect of demyelination on inhibitory neural components. We next asked whether this translated into a functional impairment of neuronal circuitry. To determine the functional consequences of these structural changes, we performed whole-cell patch-clamp recordings from layer 2 neurons within the LPC-induced demyelinated lesion in the motor cortex compared to PBS controls at 2 weeks post-surgery. To assess the formation of functional synapses, we recorded miniature excitatory and inhibitory postsynaptic currents (mEPSC and mIPSC) recorded at −70 mV and 0 mV, respectively, in the presence of 500 nM tetrodotoxin (TTX, **Fig. 4I**). We could reliably identify mEPSCs, resulting from putative AMPA receptor-mediated excitatory synapses and mIPSCs, arising from putative GABAA receptor-mediated inhibitory synapses, which reflect synapses located closest to the soma (32). The properties of mEPSCs was similar to previous reports from layer 2/3 of the motor cortex (33). Consistent with our histological analysis, we observed no difference in the amplitude **(Fig. 4L)**, frequency **(Fig. 4M)** or charge-transfer **(Fig. 4O)** of mEPSCs in layer 2 neurons from demyelinated or control animals. To assess the function of inhibitory synapses, we next assessed mIPSC properties. In neurons from demyelinated lesions at 2 weeks post-surgery, there was a significant 25% decrease in mIPSC amplitude **(Fig. 4L)**, which was accompanied by a 32% decrease in mIPSC frequency **(Fig. 4M)**. Together, this resulted in a net 44% reduction in total inhibitory charge **(Fig. 4N)**, consistent with a major loss of inhibitory synapses. These data verify a functional effect, consistent with of the observed inhibitory synapse loss after demyelination.

These electrophysiological recordings confirm that focal subpial grey matter demyelination is sufficient to cause functional loss of inhibitory synapses with no change in excitation in the cortex, in line with the selective anatomical neurode-generation of the inhibitory components.

## Discussion

We have identified a distinct neurodegenerative signature in the secondary progressive MS motor cortex characterized by the selective loss of interneuron subtypes and their connections. In a preclinical mouse model, we showed that this selective vulnerability is secondary to cortical demyelination and affects PV+, fast-spiking interneurons. The anatomical loss of PV+ interneurons is quickly translated into a functional neurophysiological impairment of the inhibitory circuitry, providing a direct link between grey matter demyelination and selective neurodegeneration of inhibitory neuronal components.

The selective effect on the inhibitory network in the human progressive MS motor cortex opposes the initial hypothesis that MS-induced neurodegeneration affects all neuronal types equally. Although at post-mortem we can only study the end-stage of a chronic and progressive disease, our results indicate that the inhibitory neural network is more susceptible to damage and/or less receptive to repair than the excitatory one. However, post-mortem analysis cannot explain the exact role of demyelination in this process as there is no temporal data. Cortical areas that appear normally myelinated may not be, either due to remyelination (which is difficult to distinguish due to sparse cortical myelination) or as they may reside next to a demyelinating lesion (excluded from the sampled section due to tissue size). Therefore, to understand if demyelination was the cause of the selective neurodegeneration in human motor cortex, the development of our mouse model was necessary.

Our preclinical model permitted the generation of a focal subpial cortical lesion of controlled and limited size that avoids the confounding effects of extensive white and grey matter demyelination or of broad cortical inflammation. A single focal demyelinated lesion in the superficial cortical layers of mouse replicated the loss of interneuron somata, axons and their inhibitory synapses, as observed in the human tissue and resulted in a functional impairment of inhibitory transmission. PV+ interneurons, PV+ neurites and inhibitory synapses were very quickly lost after demyelination which led us to the question as to why inhibitory interneurons are more at risk than pyramidal neurons after cortical demyelination. Part of the answer may rely on the location of myelinated neurons, the amount of myelination per axon and the importance of myelination for their metabolic support.

Excitatory pyramidal neurons have their soma in a specific cortical layer and send their axons over long distances, often innervating distant brain areas. Many also show discontinuous patterns of myelination (18). Conversely, most cortical interneurons have short but elaborate axonal processes that reside in the same cortical area (34). A large fraction of cortical interneuron axons are myelinated in rodents and humans, the majority of which belong to the PV+ basket cells, the most abundant cortical interneuron subtype (19–22, 35). In mouse, approximately 65% of each PV+ axon is covered by myelin compared to approximately 25% for SST+ and 10% for vasoactive intestinal polypeptide expressing interneurons suggesting that these different myelination patterns reflect nuances in the importance of myelination for each neuronal subtype (19). Heavily myelinated PV+ basket cells directly target the soma and proximal dendrites of pyramidal neurons and other interneurons mediating fast rhythmic inhibition, which requires a constant supply of ATP as manifested by their increased numbers of axonal and pre-synaptic mitochondria and cytochrome c oxidase (36, 37). This implies that PV+ axons are more reliant on the metabolic coupling between the axon and the oligodendrocyte for nutrient supply. Indeed, previous studies in which the cytochrome c oxidase pathway was ablated showed preferential deficits in the recruitment of inhibition suggesting a causal link (38). Given the location of these cells, their locally projecting axons and their dependence on myelination it is reasonable to speculate that the relative amount of myelin lost per axon is higher for PV+ interneuron axons than pyramidal ones, which renders them more susceptible to irreversible damage when the superficial cortical layers are demyelinated.

In the mouse model, the fast loss of PV+ interneurons after demyelination resulted in a marked reduction in the frequency, amplitude and charge transfer of inhibitory post-synaptic currents albeit with no effect on excitatory post synaptic currents. This functional impairment of inhibitory transmission observed in rodents is of relevance to human MS. In the healthy human motor cortex, GABAergic neurotransmission is related to motor learning and use-dependent plasticity (39, 40). In secondary progressive MS, GABA levels were found reduced in the somatosensory cortex of patients and were linked to reduced motor performance (11, 41). We now provide evidence that this selective neurodegeneration observed in both our rodent model and human MS can derive directly from cortical demyelination, primarily through loss of PV+ interneurons, their axons and synapses. This new understanding of the pattern of cortical neurodegeneration in MS allows us to think more strategically about neuroprotective therapies, perhaps by providing increased and targeted support to PV+ interneurons and selectively increasing their remyelination. Our new preclinical mouse model gives us a tractable relevant system to carry out drug screens to test this strategy with the aim of developing new treatments for progressive MS.

## Methods

### Human tissue

Post-mortem brain tissue (motor cortices) from MS patients and non-neurological controls were provided by a UK prospective donor scheme with full ethical approval from the UK Multiple Sclerosis Society Tissue Bank (MREC/02/2/39) and from the MRC-Edinburgh Brain Bank (16/ES/0084). MS diagnosis was confirmed by neuropathological means by F. Roncaroli (Imperial College London) and Prof. Colin Smith (Centre for Clinical Brain Sciences, Centre for Comparative Pathology, Edinburgh) and clinical history was provided by R. Nicholas (Imperial College London) and Prof. Colin Smith. Tables 1 and 2 include details on samples used for histological analysis and array tomography respectively. For histological analysis, tissue blocks of 2 cm x 2 cm x 1 cm were collected, fixed, dehydrated and embedded in paraffin blocks. 4 µm sequential sections were cut and stored at room temperature. Grey matter MS lesions were identified using anti-Proteolipid Protein (PLP) immunostaining. For array tomography, 1cm^3^ motor cortex blocks were collected upon autopsy and further dissected in 1mm x 1mm x 5mm array tomography samples before processing.

**Table 1.** Human case information (paraffin blocks)

| Sample ID | Age (years) | Sex | Cause of Death | Post-mortem interval (hours) | MS disease duration (years) |
| --- | --- | --- | --- | --- | --- |
| CO40 | 61 | F | Ovarian cancer | N/A | N/A |
| CO41 | 66 | M | Carcinoma of lung | N/A | N/A |
| CO53 | 89 | F | Hypertension, Type II diabetes, Hyperthyroidism | N/A | N/A |
| CO59 | 86 | M | Myocardial infarction | 38 | N/A |
| CO67 | 67 | F | Metastatic Ovarian Cancer | 32 | N/A |
| CO72 | 77 | M | Pneumonia, Ischaemic bowel | 26 | N/A |
| CO73 | 71 | M | Liver Cancer | 29 | N/A |
| CO74 | 84 | F | Old age | 22 | N/A |
| CO75 | 88 | M | COPD | 8 | N/A |
| CO76 | 87 | M | Pneumonia, Idiopathic pulmonary fibrosis | 31 | N/A |
| BBN_24780 | 57 | F | Coronary artery atheroma | 113 | N/A |
| BBN_14395 | 74 | F | Pulmonary thromboembolism | 41 | N/A |
| BBN_3778 | 70 | M | Hypertensive + ischaemic heart disease; type 2 diabetes; fatty liver degeneration. | 74 | N/A |
| BBN_18391 | 58 | M | Coronary artery thrombosis; coronary artery atherosclerosis. | 49 | N/A |
| BBN001.29693 | 49 | F | Found dead | 94 | N/A |
| BBN001.29880 | 57 | M | Found dead in bed | 64 | N/A |
| MS236 | 57 | F | Pneumonia, SPMS | 4 | 22 |
| MS239 | 69 | F | Bronchopneumonia, SPMS | 42 | 51 |
| MS243 | 61 | M | Sigmoid volvulus, SPMS | 77 | 40 |
| MS246 | 55 | M | Ischaemic heart disease, coronary atheroma, SPMS | 69 | 24 |
| MS250 | 60 | F | Bronchopneumonia, SPMS | 38 | 32 |
| MS285 | 61 | F | Paracetamol overdose, SPMS | 108 | 21 |
| MS287 | 55 | M | SPMS | 73 | 31 |
| MS290 | 66 | M | Bronchopneumonia, SPMS | 38 | 51 |
| MS309 | 74 | M | Bronchopneumonia, SPMS | 23 | 37 |
| MS343 | 71 | F | Septicaemia, bronchopneumonia, SPMS | 23 | 35 |
| MS344 | 57 | F | Septicaemia caused by urinary tract infection, SPMS | 14 | 15 |
| MS355 | 59 | F | Pneumonia, multiple sclerosis, SPMS | 8 | 52 |
| MS357 | 80 | F | Carcinomatosis, breast cancer, SPMS | 27 | 32 |
| MS359 | 52 | F | Pneumonia | 32 | 17 |
| MS361 | 60 | F | Advanced sigmoid colon cancer, SPMS | 10 | 34 |
| MS365 | 75 | F | Left lower lobe pneumonia, SPMS | 27 | 36 |
| MS385 | 64 | F | Aspiration pneumonia, SPMS, atrial fibrillation | 47 | 40 |
| MS391 | 85 | F | Aspiration pneumonia | 24 | 54 |
| MS392 | 74 | F | Bladder cancer, SPMS | 19 | 41 |
| MS417 | 74 | M | Myocardial Ischaemia, coronary artery thrombosis, coronary atherosclerosis, Infarcted small intestine within hernia, SPMS | 19 | 48 |
| MS429 | 50 | M | Sepsis, intussusception, SPMS | 42 | 24 |
| MS433 | 57 | F | Pneumonia, SPMS | 12 | 35 |
| MS503 | 53 | F | Bronchopneumonia, osteomyelitis, Acute pancreatitis, Gallstone obstruction, SPMS | 34 | 30 |
| MS507 | 67 | M | Chest infection, SPMS, Diabetes Mellitus | 42 | 40 |
| MS510 | 38 | F | Pneumonia, SPMS | 19 | 38 |
| MS515 | 72 | M | Bronchopneumonia, SPMS | 12 | 21 |
| MS531 | 58 | F | SPMS | 30 | 22 |
| MS544 | 73 | F | SPMS | 18 | 39 |
| MS549 | 50 | M | End stage SPMS | 8 | 29 |
| MS567 | 45 | F | SPMS | 48 | 23 |
| BBN_2317 | 50 | M | SPMS, Combined effects of hypertensive heart disease/cirrhosis; alcoholism; heart disease - other. | 47 | 14 |
| BBN_2391 | 57 | F | MS; Bronchopneumonia. | 7 | N/A |
| BBN_2417 | 70 | M | Ischaemic heart disease; Coronary artery atheroma; Chronic obstructive pulmonary disease; MS. | 45 | N/A |
| BBN_22229 | 65 | F | C. Diff/Sepsis. MS | 50 | NA |
| MS242-11 | 57 | F | MS | 12 | 19 |

**Table 2.** Human case information for AT tissue.

| Sample ID | Age (years) | Sex | Cause of Death | Post mortem Interval (hours) | MS disease duration (years) |
| --- | --- | --- | --- | --- | --- |
| BBN001.29529 | 42 | M | Suicide | 103 | N/A |
| BBN001.29531 | 42 | M | Sudden collapse while cycling | 94 | N/A |
| BBN001.29540 | 50 | M | Sudden collapse in street | 70 | N/A |
| BBN001.29533 | 57 | M | Coronary artery atheroma | 110 | N/A |
| BBN001.29693 | 49 | F | Found dead | 94 | N/A |
| BBN001.29731 | 53 | M | Suicide | 96 | N/A |
| BBN001.29906 | 51 | M | Sudden collapse<br>Haemopericardium | 52 | N/A |
| BBN001.30140 | 50 | M | Dissection of thoracic aorta | 122 | N/A |
| BBN001.29880 | 57 | M | Found dead in bed | 64 | N/A |
| BBN001.30169 | 48 | M |  |  | N/A |
| BBN001.32331 | 55 | M | Mesenteric ischaemia, SPMS |  | 20 |
| BBN001.32848 | 61 | F | Multi-organ failure secondary to sepsis, SPMS | 80 | 25 |
| BBN001.30890 | 70 | F | SPMS | 67 | 21 |
| BBN001.32819 | 70 | F | Mid brain stroke, Cerebrovascular disease, Type 2 diabetes, Hypertension, SPMS | 48 | NA |
| MS636 | 62 | M | SPMS | 31 | 18 |
| MS640 | 69 | F | aspiration pneumonia and SPMS | 26 | 40 |
| MS641 | 69 | F | COD bronchopneumonia and SPMS | 24 | 19 |
| MS644 | 69 | F | SPMS, bronchopneumonia, osteoporosis, | 15 | 22 |
| MS645 | 83 | F | End stage SPMS | 45 | 38 |
| MS646 | 47 | F | Aspiration, MS (RR) | 9 | 17 |
| MS658 | 69 | F | ischaemic bowel, small bowel obstruction and SPMS | 38 | 40 |
| MS665 | 74 | F | pneumonia and end stage SPMS | NA | 50 |
| MS669 | 84 | F | frailty of old age and SPMS | 12 | 59 |
| MS672 (or BBN001.35096) | 64 | F | Colon Cancer and SPMS | 44 | 31 |
| MS691 | 65 | F | Bronchopneumonia and SPMS | NA | 25 |

**Table 3.** Antibody information.

| Antibodies | Source | Identifier | Dilution |
| --- | --- | --- | --- |
| Monoclonal IgG2b anti-Islet-1&Islet-2 homeobox | Developmental Studies Hybridoma Bank | Cat# 39.4D5, RRID:AB_2314683 | 1/50 |
| Polyclonal anti- GAD65 & GAD67 | Abcam | Cat# ab11070, RRID:AB_297722 | 1/150 |
| Polyclonal anti-parvalbumin | Swant | Cat# PV27, RRID:AB_2631173 | 1/1000 |
| Monoclonal IgG1 anti-Neurofilament Marker (pan axonal, cocktail) | Biolegend | Clone SMI 312, Cat# 837904, RRID:AB_2566782 | 1/100 |
| Monoclonal IgG2b anti-Somatostatin | MERCK-Millipore | clone YC7, Cat# MAB354, RRID:AB_2255365 | 1/250 |
| Monoclonal anti-Calretinin (CR) | Swant | Cat# 6B3, RRID:AB_10000320 | 1/100 |
| Polyclonal anti-Cholecystokinin | Abcam | ab134713 RRID: N/A | 1/100 |
| monoclonal IgG1 anti Calbindin D-28k antibody | Swant | Cat# 300, RRID:AB_10000347 | 1/100 |
| Polyclonal Anti-Synapsin I | MERCK-Millipore | Cat# AB1543P, RRID:AB_90757 | 1/250 |
| Monoclonal IgG2a, anti-PSD-95 MAGUK scaffolding protein | UC Davis/NIH NeuroMab Facility | Clone: k28/43, Cat# 75-028, RRID:AB_2292909 | 1/250 |
| Polyclonal anti-Vesicular Glutamate Transporter 1 | MERCK-Millipore | Cat# AB5905, RRID:AB_2301751 | 1/250 |
| Polyclonal anti-Vesicular Glutamate Transporter 2 | MERCK-Millipore | Cat# AB2251-I, RRID:AB_2665454 | 1/250 |
| Polyclonal anti- Anti-VGAT (GABA transporter in the membrane of synaptic vesicles) | Synaptic Systems | Cat# 131 004, RRID:AB_887873 | 1/250 |
| Monoclonal IgG1 anti-gephyrin | Synaptic Systems | Clone: mAb7a, Cat# 147 011C3, RRID:AB_887716 | 1/100 |
| Monoclonal IgG2a anti-MBP (aa82-87) | Bio-Rad | Cat# MCA409S, RRID:AB_325004 | 1/300 |
| Monoclonal IgG anti Iba1 | Abcam | Clone: EPR16588, Cat# ab178846, RRID:AB_2636859 | 1/500 |
| Polyclonal anti- Glial Fibrillary Acidic Protein | Agilent | Cat# Z0334, RRID:AB_10013382 | 1/500 |
| Polyclonal anti-CD3 | abcam | Cat# ab5690, RRID:AB_305055 | 1/100 |

### Animals

Mice were housed and used according to the standard UK Home Office regulations, under project license PADF15B79. PSD95-eGFP animals were a gift from S.G.N.Grant (Centre for Clinical Brain Sciences, University of Edinburgh) (42). 8-10 week old male mice were used for all experiments.

### Array tomography (AT)

Dissected tissue blocks were processed as previously described (30). Briefly, small tissue blocks containing all 6 cortical layers were fixed for 2-3 hours in 4% PFA, 2.5% sucrose in 0.01M PBS and washed in 3.5% sucrose, 50mM glycine in 0.01mM PBS at 4°C overnight. AT samples were then dehydrated in graded series of ethanol for 5 minutes each, followed by 5 minute incubation with 50:50 ethanol 100% LRWhite (London Resin Medium grade, Agar scientific) and 100% LRWhite alone. Samples were then left overnight at 4°C in LRWhite for complete infiltration, transferred to gelatin capsules filled with cold LRWhite, and polymerized at 52°C overnight. Tissue was sectioned into ribbons of 70nm serial sections (30-40 sections/ribbon) with an ultramicrotome (Leica Ultracut) using a Ultra Jumbo Diamond Knife 35° (Diatome) and mounted in gelatin covered coverslips that were allowed to air dry before immunohistochemistry.

### LPC-induced cortical demyelinating lesion in cortex

In 8-10 week old C57BL/6 or PSD95-eGFP anesthetized male mice; a craniotomy was made above the left motor cortex. Briefly, using a 0.6mm dental drill tip and low drilling speed we thinned the skull above the M1 and M2 areas (2mm Ø) avoiding brain overheating. When the skull was adequately thinned saline application allows for the removal of the cranial top avoiding damage to the underlying brain tissue. Macroporous, poly(ethylene glycol) (PEG) based cylindrical scaffolds (cryogel) were synthesized as previously reported (31) to dimensions 2mm Ø x 0.5 mm depth and loaded with either PBS (control) or 10mg/mL L-α-Lysophosphatidylcholine (LPC, L4129, Sigma-Aldrich) and placed onto the exposed cortical surface. The skin was subsequently sutured back over the cryogel and the animals were left to recover. Two time points were chosen for the present study (2 and 3 weeks post-surgery), characterized by the presence of focal demyelination in the superficial layers of the motor cortex. Mice were subsequently perfused with 4% PFA (Sigma-Aldrich), the brain tissue was harvested, cryoprotected in 30% sucrose, frozen in 2-methyl-butane (Sigma-Aldrich) and stored at −80°C.

### Immunohistochemistry

#### Human post-mortem brain tissue

Paraffin sections were rehydrated and microwaved for 15 minutes in Vector Unmasking Solution for antigen retrieval (H-3300, Vector). For chromogenic immunohistochemistry, endogenous peroxidase and alkaline phosphatase activities were blocked for 10 minutes with Bioxal solution (SP-6000, Vector). Sections were then blocked with 2.5% normal horse serum (S-2012, Vector) for 1 hour at room temperature. Primary antibodies were incubated in antibody diluent solution (003118, Thermo Fisher Scientific), overnight at 4°C in a humidified chamber. The next day, horse peroxidase or alkaline phosphatase-conjugated secondary antibodies (Vector) were applied for an hour at room temperature. Staining was developed with either DAB peroxidase substrate kit or alkaline phosphatase substrate kit (both from Vector) according to the manufacturer’s instructions. For immunofluorescence, sections were incubated with Autofluorescent Eliminator Reagent (2160, MERCK-Millipore) for 1 minute and briefly washed in 70% ethanol after antigen retrieval. The sections were subsequently incubated with Image-iT® FX Signal Enhancer (I36933, Thermo Fisher Scientific) for 30 minutes at room temperature, washed and blocked for 1 hour with 10% normal horse serum, 0.3% Triton-X in PBS. Primary antibodies were diluted in antibody diluent solution (as above), incubated overnight at 4°C in a humidified chamber. The next day sections were incubated with Alexa Fluor secondary antibodies (Thermo Fischer Scientific, 1:1000) for 1 hour at room temperature and counterstained with Hoechst for nuclear visualization. All slides were mounted using Mowiol mounting medium (475904, MERCK-Millipore).

#### Array tomography

Dried ribbons were incubated in 50mM glycine in 0.01 M PBS for 5 minutes, washed in 3.5% sucrose, 50mM glycine in 0.01mM PBS and blocked with 0.1% BSA, 0.05% Tween-20 in Tris Buffered Saline solution (TBS) for 30 minutes at room temperature. Primary antibodies were diluted in blocking solution (all antibodies 1:50 dilution) and placed on ribbons overnight at 4°C. The following day ribbons were washed in TBS and incubated with secondary antibodies diluted in blocking solution with 0.01mg/mL DAPI for 30 minutes at room temperature (all secondary antibodies diluted 1:100). Finally, ribbons are washed and mounted on microscope slides with Shandon Immunomount (Thermo Scientific).

#### Mouse tissue

10µm thick cryosections were briefly washed in PBS and microwaved for 15 minutes in Vector Unmasking Solution for antigen retrieval (H-3300, Vector) before blocking with 10% normal horse serum, 0.3% Triton-X in 1xPBS for 1 hr at room temperature. Primary antibodies were diluted in antibody diluent (003118, Thermo Fisher Scientific) and sections were incubated overnight at 4°C in a humidified chamber. The following day, cryosections were incubated with Alexa Fluor secondary antibodies (Thermo Fischer Scientific, 1:1000) for 11/2 hrs at room temperature and counterstained with Hoechst for nuclear visualization. All slides were mounted using Mowiol mounting medium (475904, MERCK-Millipore).

### Image acquisition and analysis

#### Human post-mortem tissue

Entire sections were imaged using the Zeiss AxioScan Slide scanner and all quantifications were performed using Zeiss Zen lite imaging software.

For cell density quantification 5-10 fields were chosen that included all cortical layers, manually quantified and presented as cells/mm^2^. GM thickness was manually quantified by measuring the distance from the cortical surface to the lower edge of cortical layer six. At least 20 different measurements were obtained per section. Axonal measurements were obtained after double blinded quantification of SMI312-possitive axons as previously described (43). Briefly, the relative axonal density crossing 100µm line was measured from 10-20 different areas depending on the section size.

For synaptic density quantification, 5-6 stacks (184.58µm x 184.58µm each) from layers 2/3 of the motor cortex were obtained using high resolution confocal microscopy (Leica TCS SP8; 3144×3144 resolution, 150nm optical z-step, 2 micrometers total thickness). Each field was subdivided in 10µm x 10µm regions of interest covering the area of the neuropil avoiding cell bodies and blood vessels. Images were cropped and segmented using automated local thresholding Fiji algorithms. Segmentation parameters were exclusive for each channel but the same for all sections. To avoid false positive signal, objects that were not present in at least two consecutive sections were removed. Quantification of adjacent pre and post synaptic objects (object centers within 1µm distance) in the tissue volume was obtained using an in-house MATLAB algorithm. Values were averaged for each animal and presented as synapses/mm^3^.

#### Array tomography

Images were obtained using a Zeiss AxioImager Z2 epifluorescent microscope with a CoolSnap digital camera and AxioImager software with array tomography macros (Carl Zeiss, Ltd, Cambridge UK). A tile scan of the ribbon is initially taken in low magnification followed by the generation of a position list that outlines the area of the ribbon. Serial high-resolution images are then taken with a 63×1.4 NA Plan Apochromat objective and aligned using MultiStackReg 1.4 ImageJ plugin (30). Two to five image stacks were captured per tissue block per case. Synaptic quantification was performed as described above.

#### Mouse tissue

All sections were imaged using a Leica TCS SP8 confocal microscope. For neuronal density quantifications, 2-4 fields (581.82µm x 581.82µm each) were obtained for each section. 3-5 sections were analyzed per animal and numbers were averaged for each animal. Total number of neurons was counted in the upper cortical areas (neurons/mm^2^). For glial cell density quantification 3-6 fields (150µm x150µm each) were obtained from each section depending on the lesion size. Total number of cells was quantified in the lesion and perilesion area and represented as cells/mm^2^. Perilesion area was defined as the area spanning 150µm from the lesion border. 3-5 sections were analyzed per animal and the numbers were averaged for each animal. For signal area quantification, the same size and number of areas were thresholded and automatically quantified using in-house Fiji macros. Synaptic density quantification was done similarly to human tissue. 3 stacks (184.58µm x 184.58µm each) from layers 2/3 of the motor cortex were obtained using high resolution confocal microscopy (3144×3144 resolution, 150nm optical z-step, 2 micrometers total thickness). 3-5 sections were analyzed per animal. Quantification of adjacent pre and post synaptic objects (object centers within 0.5µm distance) in the tissue volume was obtained using an in-house MATLAB algorithm. Values were averaged for each animal and presented as synapses/mm^3^. Images were randomised using the File randomizer Fiji macro prior to analysis.

### In vitro slice electrophysiology

Slices containing LPC lesion and PBS control neurons were prepared from mice which had been implanted with cryogels overlaying the motor cortex (M1), as previously described (38, 44). Briefly, mice were sedated with isofluorane, decapitated, and their brains rapidly removed and placed in ice-cold sucrose-modified artificial cerebrospinal fluid (sucrose-ACSF; in mM: 87 NaCl, 2.5 KCl, 25 NaHCO_3_, 1.25 NaH_2_PO_4_, 25 glucose, 75 sucrose, 7 MgCl_2_, 0.5 CaCl_2_); saturated with carbogen (95% O_2_/5% CO_2_). 300µm coronal brain slices were then sliced on a vibratome (VT1200S, Leica, Germany) covering the region that was overlain by the cryogel, as assessed by eye during dissections, then transferred to submerged holding chambers filled with sucrose-ACSF at 35°C for 30 min, then room temperature until needed.

Whole-cell patch-clamp recordings were performed to record miniature EPSCs and IPSCs. Slices were transferred to a submerged recording chamber which was perfused with ACSF (in mM: 125 NaCl, 2.5 KCl, 25 NaHCO_3_, 1.25 NaH_2_PO_4_, 25 glucose, 1 MgCl_2_, 2 CaCl_2_, containing 500 nM tetrodotoxin, TTX) and bubbled with carbogen, at a rate of 4-6 mL.min-1, recordings were maintained at 30± 1 °C with an inline Peltier heating device (Scientifica, Brighton, UK). Slices were first visualised under low-power magnification (4x Objective 0.2 NA, Neoplan, Olympus, Japan) with infrared differential-inference contrast microscopy using a digital camera (Sci-CamPro, Scientifica, UK) mounted on an upright microscope (SliceScope, Scientifica, UK) to the dorsal pole of the slice. Neurons were identified for recording with a high-power 40x water-immersion objective lens (1.0 N.A., Olympus, Japan) and chosen based on having an ovoid somata located in the upper region of layer 2, with an apical dendrite oriented towards the pial surface. Recording pipettes were pulled from borosilicate glass capillaries (1.7mm outer/1mm inner diameter, Harvard Apparatus, UK) on a horizontal electrode puller (P-97, Sutter Instruments, CA, USA) and filled with a Cs-gluconate based intracellular solution (in mM 140 Cs-gluconate, 3 CsCl, 0.2 EGTA, 10 HEPES, 2 Na_2_ATP, 2 MgATP, 0.3 Na_2_GTP, 10 Na_2_Phosphocreatine, 2.7 Biocytin, 5 QX-314.Cl, pH=7.4, 290-310 mOsm), which gave 3-5 MΩ resistance electrodes and a measured Cl^-^ reversal of −74 mV under these recording conditions. Liquid junction potential was not corrected. Cells were rejected from further recording, if they required more than 200pA current injection to maintain a −70 mV holding potential, series resistance >30 MΩ, or series resistance changed by more than 20%. All recording was performed with a Multiclamp 700B amplifier (Molecular Devices, CA, USA) and filtered online at 2 kHz with the amplifiers 4-pole Bessel Filter and digitised at 20 kHz (Digidata1550B, Molecular Devices, CA, USA)

All in the presence of 500 mM TTX, mEPSCs were recorded from −70 mV voltage clamp. 5 minutes of continuous recording were performed with series resistance measured before and after this period. Following mEPSC recordings, the membrane potential was switched to between 0 mV and 5 mV – depending on measured mEPSC reversal and 5 minutes of mIPSCs recorded. Following recording, the series resistance was measured again to confirm recording stability. Out-side out patches were then performed on recorded neurons, to seal the included biocytin within cells. A further 2-3 cells were recorded from each slice, and then slices fixed in 4% PFA in 0.1 M PB overnight, then transferred to PBS until processed for histology.

mPSCs were detected offline using a template fitting approach, whereby exemplary mini-PSCs were fit with a triexponential curve (45), with thresholds of 4-7 imposed for detection. Following signal extraction, individual traces were excluded for analysis if they failed to exceed the threshold of 3*SD of the baseline noise. EPSCs were detected as inward currents and ISPCs as outward currents. Traces were collected in pCLAMP 9 (Molecular Devices, CA, USA) and stored on a desktop computer. All analysis of electrophysiological data was performed offline using the Stimfit electrophysiological package (46). All data collection and analysis were performed blind to treatment group.

## Acknowledgements

The authors would like to thank and acknowledge the Edinburgh Brain and Tissue Bank and the MS Society UK Tissue Bank for providing tissues used in this study. The Edinburgh Brain Bank is supported by the Medical Research Council, ethical approval NHS-RECSE (16/ES/0084). We thank Prof. Seth Grant for giving us the PSD95-GFP transgenic mouse. LZ is funded by the MS Society UK Centre grant to Edinburgh. AW is also funded by the MS Society UK and the Medical Research Council. TSJ is funded by UK Dementia Research Institute which receives funding from Alzheimer’s Research UK, the Alzheimer’s Society, and the Medical Research Council and the European Research Council (ERC) under the European Union’s Horizon 2020 research and innovation programme (Grant agreement No. 681181). BN, CW and DE are funded by the Deutsche Forschungsgemeinschaft (DFG) – Project number 320041273. SAB and PK are funded by the Simons Foundation Autism Research Initiative (529085) and the Patrick Wild Centre. We would like to thank Rosie Jackson and Chris Henstridge for their help in array tomography processing and analysis and Dr Veronique Miron, Prof. David Lyons and Prof. Charles ffrench-Constant for their comments on the manuscript.

## Supplementary Figures

**Fig. S1.**
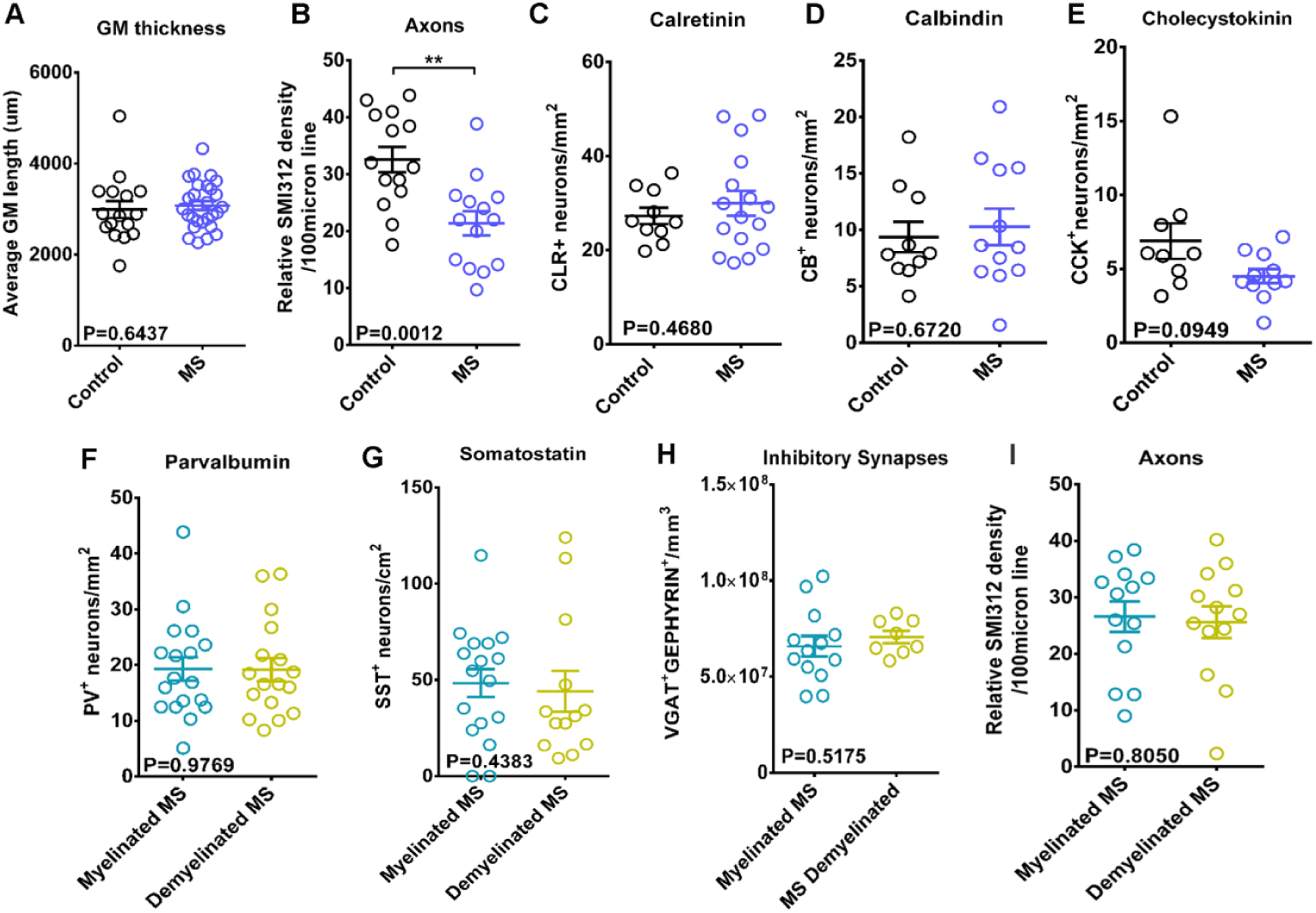
**A:** Grey matter (GM) thickness in control and MS motor cortex samples (Control: mean 2987 ± 184.6SEM µm, MS: mean 3075 ± 95.85SEM µm; each point is a patient, Unpaired t test). **B:** Relative axonal (SMI312+) density crossing 100µm line in control and MS motor cortex for layers 2/3 (Control: 32.57± 2.229 SEM number of SMI312 axons/100µm, MS: mean 21.37 ± 2.115 SEM number of axons/100µm; each point is a patient, unpaired t test). **C:** Quantification of calretinin (CR)+ interneuron density in all six cortical layers in control and MS motor cortex (Control: mean 27.27 ± 1.731 SEM neurons/mm^2^, MS: mean 29.94 ± 2.635 SEM neurons/mm^2^; each point is a patient, unpaired t test). **D:** Quantification of calbindin (CB)+ interneuron density in all six cortical layers in control and MS motor cortex (Control: mean 9.377± 1.353 SEM neurons/mm^2^, MS: mean 10.28± 1.607 SEM neurons/mm^2^; each point is a patient, unpaired t test). **E:** Quantification of cholecystokinin (CCK+) interneuron density in all six cortical layers in control and MS motor cortex (Control: mean 6.886± 1.202 SEM neurons/mm^2^, MS: mean 4.507± 0.4801 SEM neurons/mm^2^; each point is a patient, Mann Whitney test) **F:** Quantification of PV+ interneuron density in MS myelinated and demyelinated motor cortex (Myelinated MS: mean 19.27 ± 2.115 SEM neurons/mm^2^, Demyelinated MS: mean 19.19 ± 2.085 SEM neurons/mm^2^; each point is a patient, Unpaired t test). **G:** Quantification of SST+ interneuron density in MS myelinated and demyelinated motor cortex (Myelinated MS: mean 48.31 ± 7.245 SEM neurons/mm^2^, Demyelinated MS: mean 44.09 ± 10.55 SEM neurons/mm^2^; each point is a patient, Mann Whitney test). **H:** Quantification of inhibitory synapse density in MS myelinated and demyelinated motor cortex (Myelinated MS: mean 6.583e+007 ± 5.299e+006 SEM synapses/mm^3^, Demyelinated MS: mean 7.061e+007 ± 3.178e+006 SEM synapses/mm^3^; each point is a patient, unpaired t test). **I:** Relative axonal (SMI312+) density crossing 100µm line in MS myelinated and demyelinated motor cortex for layers 2/3 (Myelinated MS: 26.58 ± 2.719 SEM number of SMI312 axons/100µm, Demyelinated MS: mean 25.60 ± 2.816 SEM number of axons/100µm; each point is a patient, unpaired t test).

**Fig. S2.**
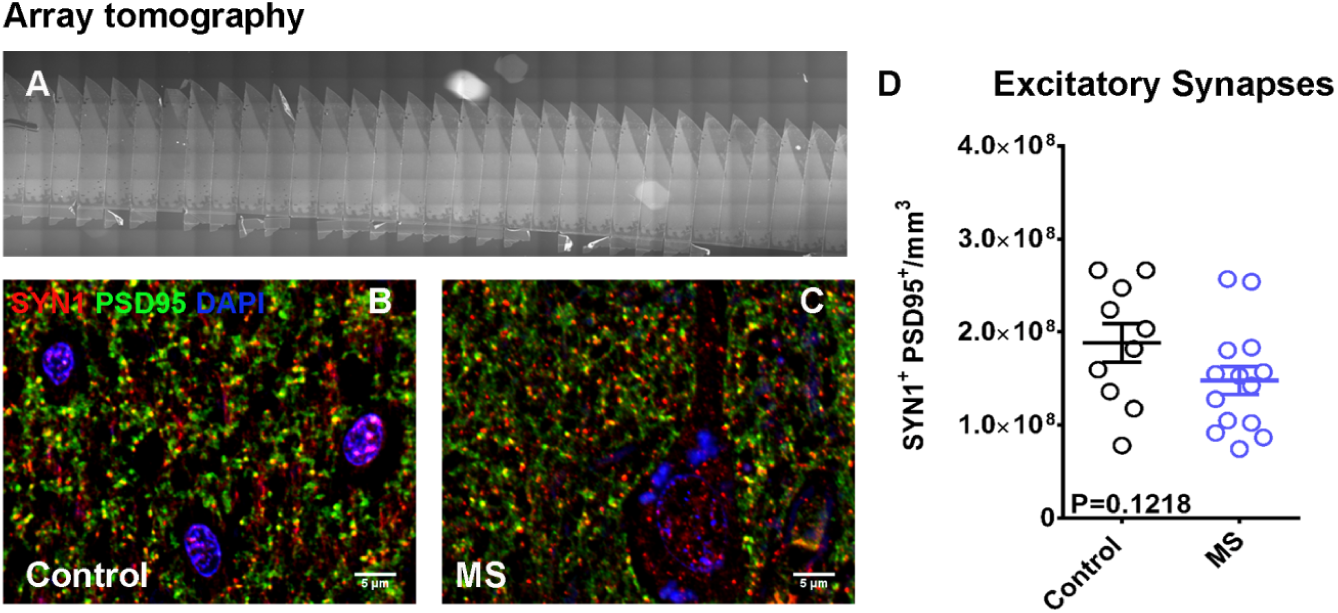
**A:** Example of an array tomography (AT) ribbon (image taken with fluorescent microscope). Each ribbon consists of sequential 70nm-thick tissue sections. **B-C:** Representative images of AT sections from control (B) and MS (C) motor cortex, stained for SYN1 (red), PSD95 (green) with DAPI (blue). Scale bar: 5µm. **D:** Quantification of excitatory, SYN1+/PSD95+ synapses of control and MS motor cortex (AT samples); (Control: mean 1.881e+008 ± 2.054e+007 SEM synapses/mm^3^, MS: mean 1.479e+008 ± 1.524e+007 SEM synapses/mm^3^; each point is a patient, unpaired t test).

**Fig. S3.**
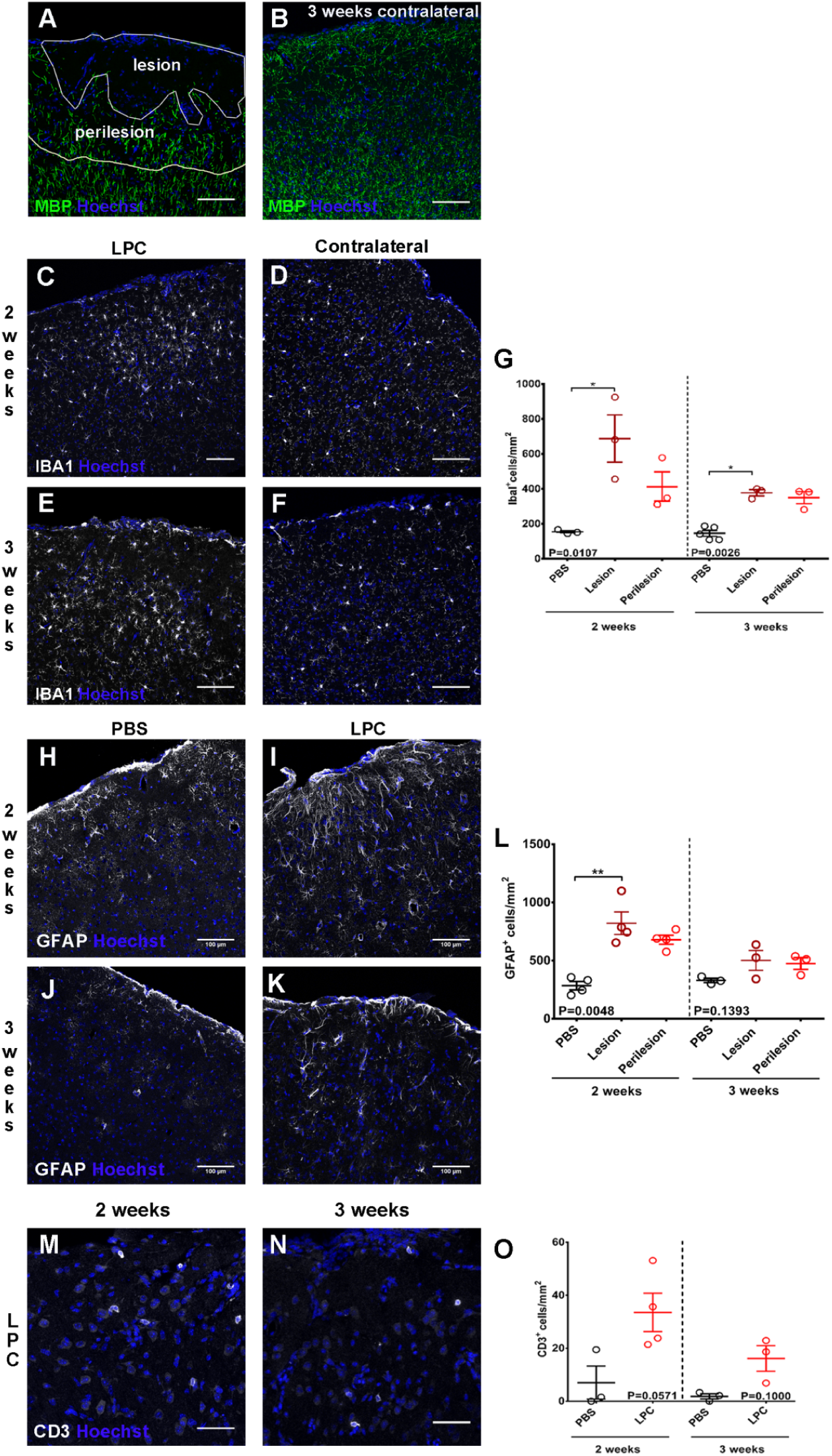
**A:** Outline of a cortical lesion and perilesion areas after LPC-loaded cryogel placement onto the rodent cortex. Cortical coronal section stained for MBP (green) and Hoechst (blue). The area 150µm from the lesion border was considered as the perilesion area. Scale bar: 100 µm. **B:** Cortical section of the contralateral hemisphere of animals treated with LPC-loaded cryogel, 3 weeks post-surgery. Immunohistochemistry for MBP (green) and Hoechst (blue). Scale bar: 100µm. **C-F:** Immunohistochemistry of LPC-loaded cryogel animals for IBA1 (white) and Hoechst (blue) two (C) and three weeks (E) after cryogel placement. Contralateral hemisphere of the 2 week (D) and 3 week (F), LPC-loaded cryogel. Scale bar: 100µm. **G:** Quantification of IBA1+ cell density at the upper cortical layers of PBS treated cortex (control) LPC-treated lesion and perilesion areas, for two (left) and three (right) weeks post –surgery (2weeks PBS: mean 154.1 ± 6.606 SEM cells/mm^2^, lesion: mean 687.3 ± 135.4 SEM cells/mm^2^, perilesion: mean 412.6 ± 83.65 SEM cells/mm^2^; 3 weeks PBS: mean: 145.5 ± 16.66 SEM cells/mm^2^, lesion: mean 377.7 ± 17.58 SEM cells/mm^2^, perilesion: mean 349.6 ± 34.08 SEM cells/mm^2^; each point is an animal, Kruskal-Wallis test with Dunn’s multiple comparisons test). **H-I:** Immunohistochemistry of PBS treated animals (H) and LPC treated (I) for GFAP (white) and Hoechst (blue) two weeks after cryogel placement. Scale bar: 100µm. **J-K:** Immunohistochemistry of PBS treated animals (J) and LPC treated (K) for GFAP (white) and Hoechst (blue) three weeks after cryogel placement. Scale bar: 100µm. **L:** Quantification of GFAP+ cell density in the upper cortical layers of PBS treated cortex (control) LPC-treated lesion and perilesion areas, two (left) and three (right) weeks post –treatment (2 weeks PBS: mean 284.9± 36.02 SEM cells/mm^2^, lesion: mean 821.9 ± 96.71 SEM cells/mm^2^, perilesion: mean 679.7 ± 39.53 SEM cells/mm^2^; 3 weeks PBS: mean 329.8 ± 18.18 cells/mm^2^, lesion: mean 502.3 ± 86.19 SEM cells/mm^2^, perilesion: mean 475.2 ± 50.20 SEM cells/mm^2^; each point is an animal, Kruskal-Wallis test with Dunn’s multiple comparisons test). **M-N:** Immunohistochemistry of LPC treated animals two (M) and three (N) weeks after cryogel placement for CD3 (white) and Hoechst (blue). In both cases the area of the lesion is shown. Scale bar: 50µm. **O:** Quantification of CD3+ cell density in the upper cortical layers of PBS treated cortex (control) LPC-treated areas, two (left) and three (right) weeks post –surgery (2 weeks PBS: mean 7.012 ± 6.246 SEM cells/mm^2^, LPC: mean 33.51 ± 7.246 SEM cells/mm^2^; 3 weeks PBS: mean 1.820 ± 0.9724 SEM cells/mm^2^, LPC: mean 16.15 ± 4.814 SEM cells/mm^2^; each point is an animal, Mann Whitney test).

## Notes

### Competing Interest Statement

The authors have declared no competing interest.

